# Ovothiol is a redundant part of a complex thiol network in promastigote *Leishmania mexicana*

**DOI:** 10.1101/2023.12.08.570742

**Authors:** Victoria L. Bolton, Clement Regnault, Lesley S. Morrison, Supriya Khanra, Ryan Ritchie, Kerrie E. Hargrave, Richard J.S. Burchmore, Megan K.L. MacLeod, Michael P. Barrett

**Affiliations:** University of Glasgsow, School of Infection, Immunity and Inflammation; Glasgow Polyomics, University of Glasgow

## Abstract

A gene encoding OvoA, a key enzyme involved in the biosynthesis of ovothiol, was excised form the genome of Leishmania mexicana promastigotes using CRISPR/cas mediated gene editing. The role of the enzyme in synthesising ovothiol was confirmed since both ovothiol A and ovothiol B were lost from the metabolome of the modified cells. The OvoA knockout line had similar growth kinetics to wild-type progenitor cells and, moreover, most of the changes in metabolism that accompanied the transition of log stage growth to stationary phase were mirrored in the KO line. Significant differences, however, were observed in the ratio of the reduced and oxidised forms of the other major low molecular weight thiols, glutathione and trypanothione, indicative of a role of these other thiols in maintaining reduced ovothiol and demonstrating an interconnected network of low molecular weight thiols in these cells. The OvoA knockout cells remained infective to macrophages where promastigotes transformed to amastigote forms in a manner similar to wild-type. The knockout line was tested for sensitivity to a range of current anti-leishmanial drugs and oxidative and nitrosative stresses. While generally the absence of ovothiol caused little or no change in sensitivity to these stress-inducing agents, enhanced sensitivity to amphotericin B was noted.

**Author summary:** Ovothiol is a low molecular weight histidine-derived thiol first described in sea urchin eggs, and later found in many organisms, including the protozoa of the order Kinetoplastida, that includes human pathogens such as the Leishmania species that cause leishmaniasis. Thiol metabolism in the Kinetoplastidae has been studied in some detail, particularly with regard to an unusual bis-glutathione, spermidine conjugate named trypanothione that takes on many of the roles performed by glutathione in most other organisms. Roles for ovothiol in Leishmania have not been previously defined, although potential roles in defence against oxidative stress have been hypothesised. A gene encoding the first enzyme of the pathway involved in ovothiol production, OvoA, was excised from the *Leishmania mexicana* genome. Its role in ovothiol synthesis was confirmed as ovothiol was absent from the mutants. Little changed, however, with respect to the phenotype of these cells, including their proliferation rate, their ability to infect macrophages or their sensitivity to a range of stress inducing agents. These included several leishmanicidal drugs, oxidative and nitrosative stresses. For amphotericin B, however, the Ovothiol lacking cells were more sensitive than wild-type indicating some role in defence against the impact of this drug.

## Introduction

Ovothiol (1-N-methyl-4-mercaptohistidine), is a low molecular weight thiol, originally found in sea urchin eggs [1–4]. Subsequently, more methyl-mercaptohistidine based molecules were found in other organisms including in protozoa of the taxonomic grouping known as the Kinetoplastida which were shown to contain ovothiol A (3-methyl-5-sulfanyl-L-histidine) [5–8]. The Kinetoplastidae include important parasitic pathogens of man including the African and American trypanosomes [9] and leishmania species [10]. The role(s) of ovothiol in these organisms has remained elusive, particularly as the parasites also contain another unusual thiol, comprising two glutathione moieties linked by a spermidine creating the well-studied molecule N^1^,N^8^-bis-glutathionylspermidine (trypanothione) [11–13]. Ovothiol was shown to be more abundant in promastigote forms of leishmania than amastigotes and its levels increased in cultured cells as proliferative promastigotes transformed to the growth-arrested metacyclic promastigote form [7] which is transmitted from the sandfly vector to their mammalian hosts. Metacyclics are taken up by macrophages and exposed to the oxidative burst including an array of free radicals, indicating a possible role in defence against these potentially toxic molecules [6], with defence against nitric oxide and other nitrosative metabolites considered more likely than scavenging H_2_O_2_ given the superiority of trypanothione in this latter function [14]. A variety of mechanisms enable the parasites to survive within macrophages [15–17], however ovothiol was reportedly absent from the replicative amastigote form of the parasite [7] leading to a suggestion that any role in survival would precede reaching this part of the life cycle.

Ovothiol is synthesised by the sulfanylation followed by methylation of histidine. A gene encoding the enzyme, termed *OvoA*, was initially discovered in *Erwinia tasmaniensis* and *Trypanosoma cruzi* [18] based on homology to another histidine sulfanylase, EgtB, from Mycobacteria [19]. OvoA undertakes both the initial sulfanylation and the later methylation reactions, but a C-S lyase reaction that cleaves the sulfoxide intermediate prior to methylation is catalysed by a separate enzyme, OvoB, the identification of which remains elusive within the kinetoplastidae. Genes related to *OvoA*, however, are found in the genomes of organisms known to produce ovothiol, including the trypanosomatids, and absent from organisms that do not synthesise this metabolite. The enzymes were named as 5-histidylcysteine sulfoxide synthases (OvoAs) [18]. Recombinant versions of the OvoA from *Erwinia tasmaniensis* and *Trypanosoma cruzi* were expressed and the enzymes characterised [18].

In order to learn more about the *in situ* roles of ovothiol in *Leishmania mexicana* we deleted the gene predicted to encode OvoA using CRISPR/cas9 technology [20] and studied the ability of the parasites to grow as promastigotes and transform between proliferative stage to stationary stage cells in culture, and their sensitivity to anti-leishmanial drugs and other stress-inducing agents. We also investigated the metabolome of the parasites to identify whether loss of *OvoA* did, indeed, correlate to loss of ovothiol in these cells and, moreover, to determine what other changes to metabolism occur as a consequence of losing ovothiol. Finally, we assessed the ability of ovothiol deficient leishmania to invade macrophages and replicate as intracellular amastigotes.

## Results

### The *OvoA* gene is readily removed from the genome of *L. mexicana* promastigotes, and knockout cells have no growth defect

Constructs containing the Blasticidin-S deaminase (c*BSD*) and Puromycin *N*-acetyltransferase (*PAC*) genes flanked from the 5’ flank of the *OvoA* gene and from the 3’ flank were transfected into *Leishmania Cas9* [20]. Post-transfection, parasites were selected in blasticidin plus puromycin and the *OvoA* transfected line was found to grow in the double selective medium 11 days post transfection. Individual cells were then cloned using limiting dilution and gDNA purified from these parasites amplified using oligonucleotides from within the *OvoA* gene to verify its absence as well as the marker genes to verify their presence. The successful double gene replacement was confirmed (Supplementary Figure 1) indicating that *OvoA* is not essential to *L. mexicana* promastigotes.

Having established non-essentiality of *OvoA* we determined its impact on the growth kinetics of *L. mexicana*, by comparing the rate of growth of Δ*OvoA L. mexicana* in comparison to WT promastigote cultures, counted every 24 hours over 7 days. As demonstrated in Figure 1 only a minor difference in growth kinetics was observed between Δ*OvoA L. mexicana* and WT cell cultures. Specifically, Δ*OvoA L. mexicana* reached a slightly lower maximum cell density after three days than the WT counterpart. Both Δ*OvoA* and WT *L. mexicana* had a logarithmic phase that lasted for 3 days followed by a stationary phase from 3-7 days, with a slow loss of cell density likely due to cell death (Figure 1).

**Figure 1:**
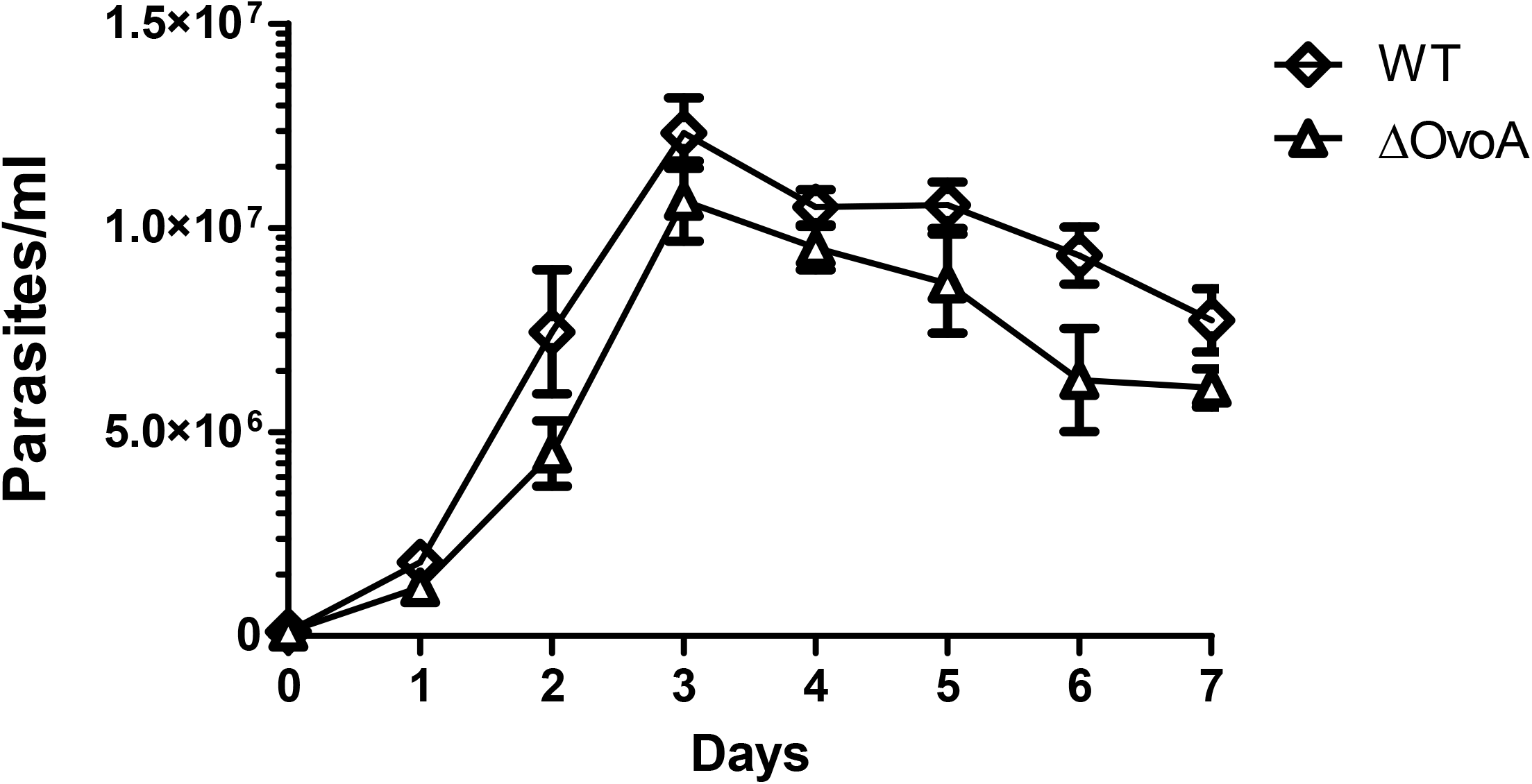
Growth kinetics of Wild type *Leishmania mexicana* and *Leishmania mexicana* Ovothiol A Knockout mutant over 7 days (n=6)

### Comparative metabolomics analysis of WT and *OvoA* knockout parasites

In order to determine whether the deletion of the *OvoA* gene leads to the absence of the Ovothiol A metabolite in the Δ*OvoA L. mexicana* line, and to ascertain any other changes associated with its loss, metabolomics analysis of metabolite extracts collected from stationary and logarithmic phase Δ*OvoA* and wildtype *L. mexicana* promastigotes was performed using LC-MS. A heatmap (supplementary figure 2) reveals the top 100 significantly changed metabolites and supplementary figures 3 and 4 respectively show pathway enrichment analysis in knockout cells at logarithmic and stationary phase respectively.

Principal component analysis (PCA) of the detected metabolites, demonstrated clear separation of the Δ*OvoA* and wildtype *L. mexicana* metabolome of both stationary and logarithmic samples along the first principal component (Figure 2A). Separation of Δ*OvoA* and wildtype *L. mexicana* groups was driven along the second principal component by the following metabolite annotations as shown in the corresponding Loadings plot (Figure 2B): ovothiol A, the peptide Gln-Thr-Gln-Tyr, ovothiol B, tetcyclasis and glutathione disulphide on the one side (Δ*OvoA*), and glutathione, trypanothione and gamma-L-glutamyl-L-cysteine-beta-alanine on the other side (WT). It is important to note that metabolite identifications are tentative in the absence of orthogonal information beyond mass, e.g. inclusion of chemical standards to match retention time or fragmentation, and in the case of tetcyclasis, which is a plant hormone for which no evidence for its existence in Leishmania exists, it is likely that the true identity of this metabolite is different. A strong separation between parasites at logarithmic and stationary-phase irrespective of their genotype was also noted in the PCA (Figure 2A).

Separation of groups in the PCA was accompanied by 93 Bonferroni corrected significant changes (adj.p-value (FDR)< 0.05) in the abundance of metabolites between logarithmic phase Δ*OvoA* and WT *L. mexicana* and 374 significantly changed metabolites (adj.p-value (FDR)< 0.05) between stationary phase Δ*OvoA* and WT *L. mexicana* (Figure 3). From these significant changes, 55 metabolites were identified and 11 metabolites were of significantly decreased abundance or increased abundance, in logarithmic phase Δ*OvoA L. mexicana* (Figure 3A), whereas 105 metabolites were significantly down- and 79 metabolites of increased abundance (threshold Log_2_FC>1) in stationary-phase Δ*OvoA L. mexicana* (Figure 3B).

**Figure 2:**
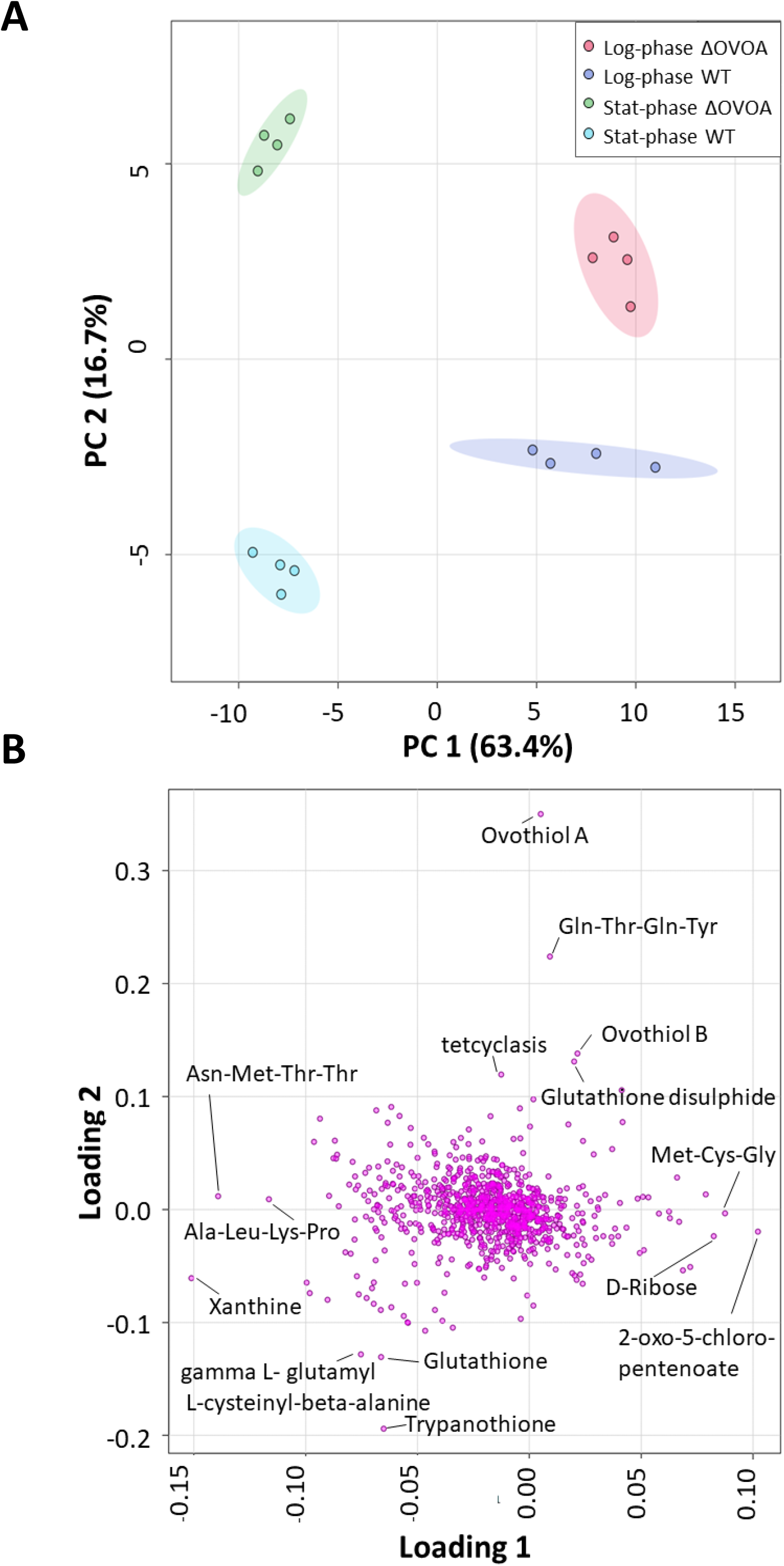
Principal Component analysis scores plot (A) and associated loading plot (B) of metabolomic samples from log-phase and stationary phase Δ*OvoA* and WT *L.mexicana*. Metabolite extracts were collected from 1x 108 log-phase or stationary-phase *OvoA* Knockout mutant and WT *L. mexicana* promastigotes. Four samples were collected per condition (n=4) and analysed on a Liquid chromatography Mass spectrometry (LC-MS). Principal Component analysis was performed with Metaboanalyst v5.0. (A) Principal component 1 (PC1) and associated variance of 63.4% is dislpayed on the X axis and Principal component 2 (PC2) and associated variance of 16.7% is dislpayed on Y axis. (B) Metabolites (loadings) that strongly influence both components are highlighted. OvoA: Ovothiol A; WT: Wild type.

**Figure 3:**
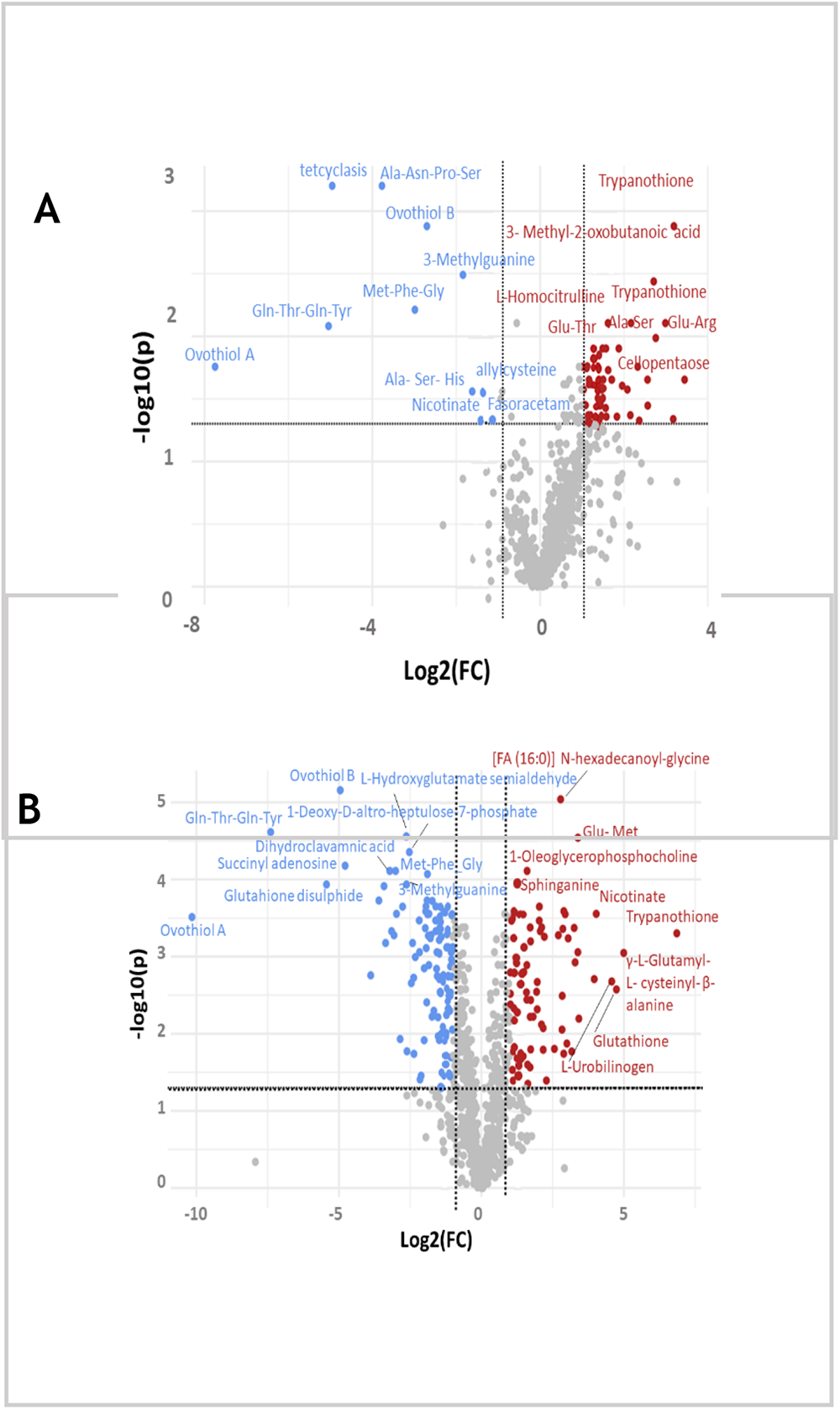
Volcano plot of metabolome from log-phase (A) and stationary-phase (B) Δ*OvoA* and Wildtype *L. mexicana.* Volcano plot displays the fold change of each metabolite in log-phase Δ*OvoA* and Wildtype *L. mexicana* against the adjusted -log10 p-values. Metabolites in red are significantly upregulated. Proteins in blue are significantly downregulated (FDR, p=<0.05, log_2_FC=>1). Figure was made with metaboanalyst vs. 5.0.

Ovothiol A and ovothiol B were absent in both stationary-phase and logarithmic-phase Δ*OvoA L. mexicana* (Figure 3) (note that the Ideom software attributes a notional AUC value of 1,000 to absent metabolites in order that a comparison, albeit artificial, is possible between datasets where a metabolite may be included in one but not the other). Figure 4, reveals the absence of both Ovothiol A and B in Δ*OvoA L. mexicana* as compared to the wildtype parasites which confirms the role of the putative *OvoA* gene in the biosynthesis of ovothiol.

**Figure 4.**
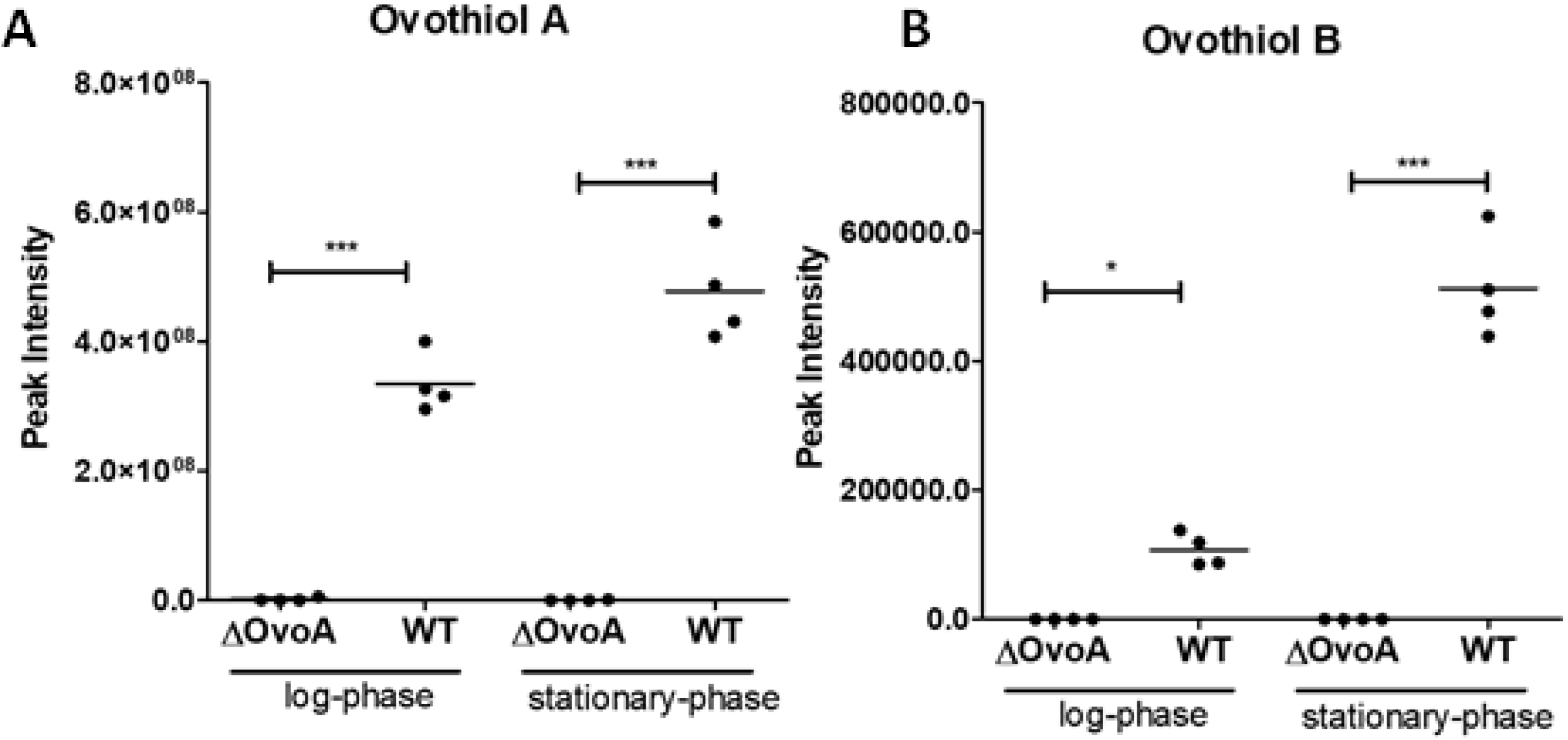
Peak intensities of ovothiol A (A) and ovothiol B (B) in logarithmic and stationary phase ΔOvoA and WT L. mexicana. Each group is composed of four metabolomic samples (n=4). Statistical significance was tested using a one-way ANOVA with Bonferroni’s Multiple Comparison Test as a post-hoc test (***=p<0.001; *=p<0.05).

Other metabolites that were diminished in abundance in both logarithmic and stationary-phase Δ*OvoA L. mexicana* were tentatively annotated as 3-methylguanine, tetcyclasis (considered an unlikely annotation as noted above) and peptides such as Gln-Thr-Gln-Tyr and Met-Phe-Gly (Figures 2 & 3). Metabolite enrichment analysis of metabolites of significantly decreased abundance in the Δ*OvoA* mutant revealed some pentose phosphate pathway-related metabolites (e.g. deoxy-D-aldro heptulose 7P) and several metabolites associated with the metabolism of alanine, aspartate and glutamine to be lowered in mutant *L. mexicana* in both logarithmic and stationary-phase (Supplementary Figure S3 & S4). Overall, fewer metabolites of significantly lowered abundance were observed in Δ*OvoA* in comparison to the WT *L. mexicana* cell line in logarithmic-phase compared to stationary-phase cells (Figure 3).

In contrast, pathways such as arginine synthesis, valine, leucine and isoleucine synthesis as well as glutathione biosynthesis were found to be increased in both logarithmic and stationary-phase ΔOVOA *L. mexicana* (Supplementary Figures S3 & S4). Reduced glutathione was significantly enriched in stationary as well as logarithmic-phase ΔOVOA *L. mexicana* (Figure 5). By contrast, the oxidised form of this thiol, glutathione disulfide, was found to be diminished in ΔOVOA *L. mexicana* (Figure 5).

Of three individual peaks corresponding to the mass of the putative metabolite trypanothione, (Supplementary Figure S6) only that with a retention time of 12.27 matched a trypanothione standard (Supplementary Figure S7), others may be fragmentation products of unidentified trypanothione adducts. The standard matched trypanothione peak significantly increased in logarithmic-phase *OvoA* KO compared to WT *L. mexicana.* A slight, non-significant, increase for this metabolite was also noted in stationary-phase *OvoA* KO, whereas trypanothione was not identified in WT stationary-phase *L.mexicana*. The oxidised form of trypanothione, trypanothione disulfide (T(S)_2_, provided a peak with a relatively strong signal that was diminished, albeit not reaching statistical significance, in Δ*OvoA L. mexicana* (Figure 6).

This is consistent with a significant proportion of the reduced forms of glutathione and trypanothione usually being used in maintaining ovothiol in a reduced form. Loss of ovothiol then alters the cellular redox balance such that trypanothione and glutathione are required to give up fewer protons than usual in order to sustain this reduced ovothiol, and hence these reduced forms increase in abundance while their oxidised counterparts are diminished. Trypanothione is the central redox thiol and maintained in its reduced form by the enzyme trypanothione reductase. Non-enzymatic reduction of glutathione by trypanothione, and ovothiol by either of these metabolites ensues (Figure 7).

**Figure 5:**
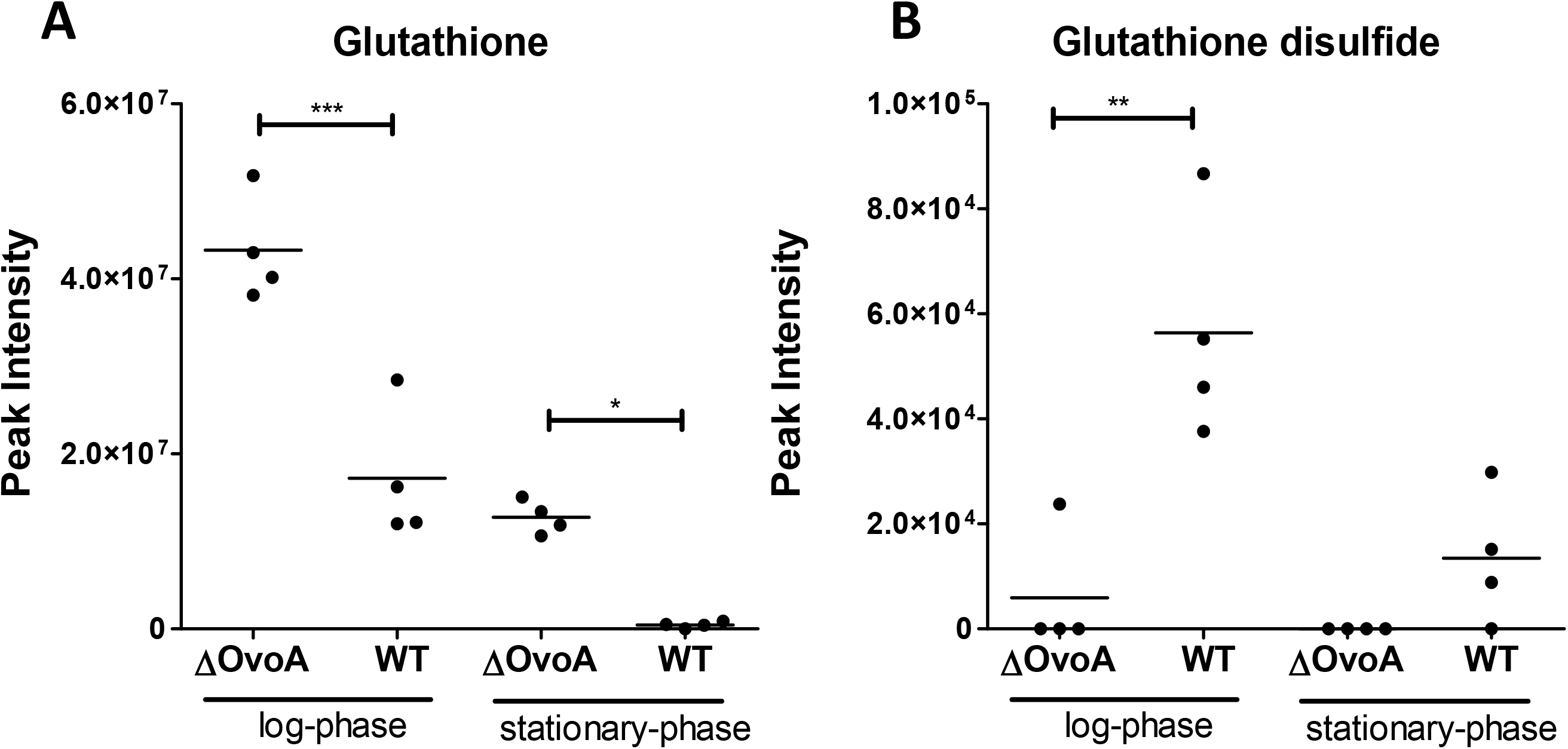
Peak intensities of putative Glutathione A (A) and Glutathione disulfide (B) in logarithmic and stationary phase Δ*OvoA* and WT *L. mexicana.* Each group is composed of four metbolomic samples (n=4). Statistical significance was tested using a One-way ANOVA with Bonferronis Multiple Comparison Test as a post-hoc test (***=p<0.001; **=p<0.01*=p<0.05).

**Figure 6:**
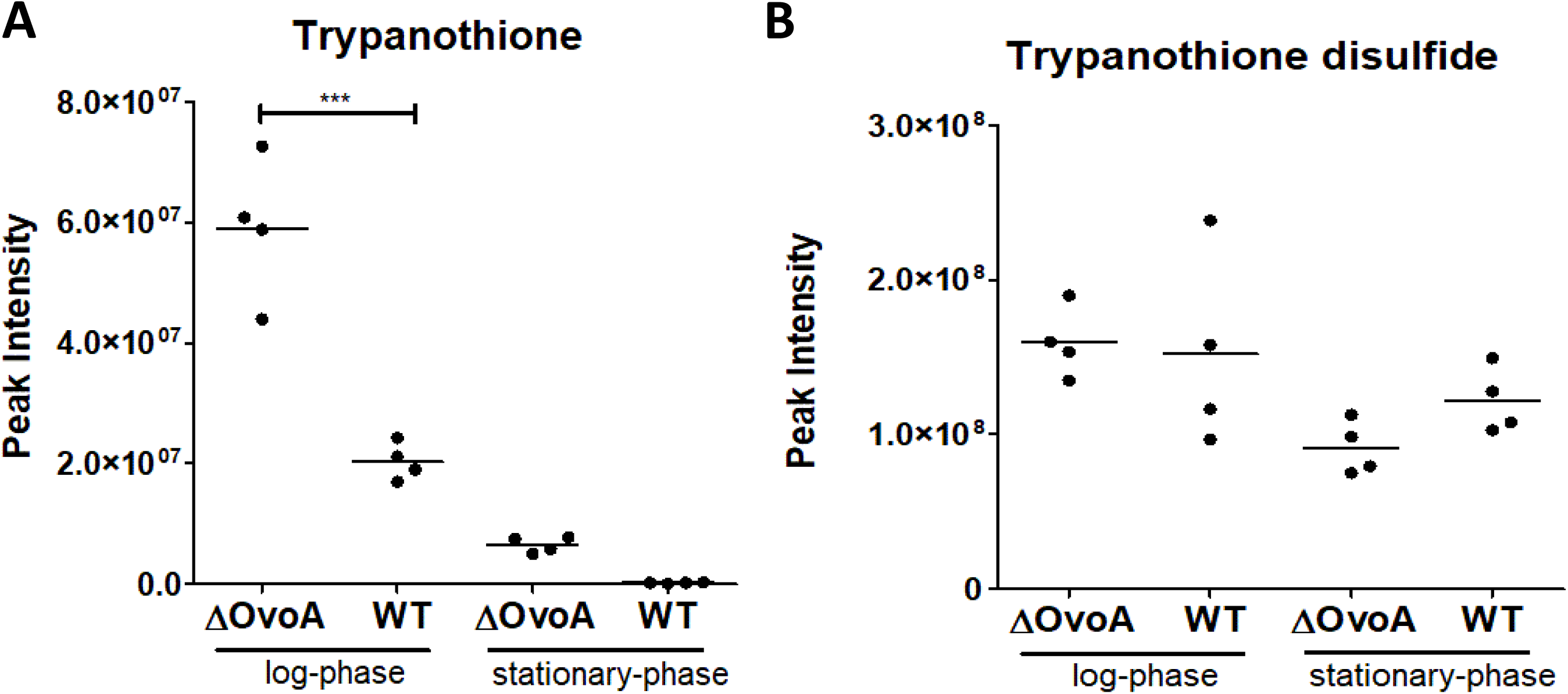
Peak intensitiy of trypanothione peak matching to a trypanothione authentic standard and (A) and of a putative trypanothione disulfide (B) extracted from ideome in logarithmic and stationary phase Δ*OvoA* and WT *L. mexicana.* Each group is composed of four metbolomic samples (n=4). Statistical significance was tested using a One-way ANOVA with Bonferronis Multiple Comparison Test as a post-hoc test (***=p<0.001; *=p<0.05).

Other metabolites that were increased in ΔOVA *L. mexicana* were the hexose polymer metabolites labelled as cellopentaose and cellohexose (Figure 3). These likely represent the five and six chain representatives of poly-mannose, part of the key leishmania storage carbohydrate, mannogen [21].

**Figure 7:**
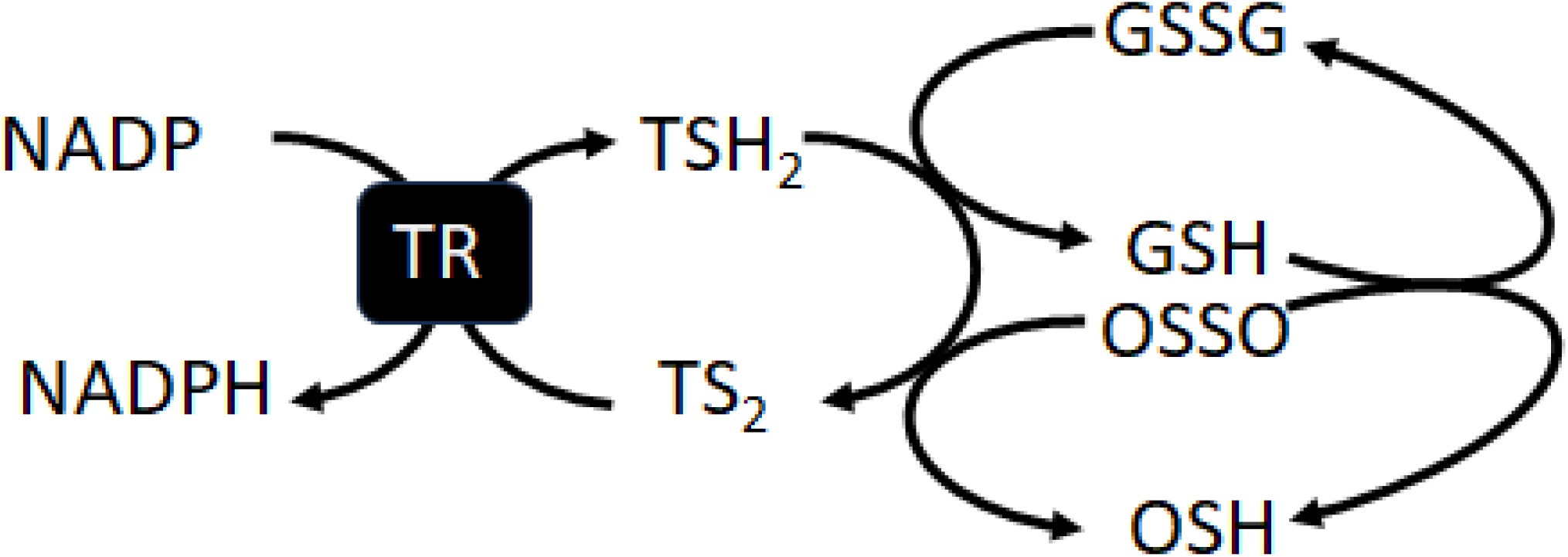
Scheme of the low molecular weight thiol network in *Leishmania mexicana*. The enzyme trypanothione reductase (TR) reduces trypanothione disulfide (TS_2_) using protons from NADPH. Reduced trypanothione (TSH_2_) can then directly reduce glutathione disulfide (GSSG) to reduced glutathione (GSH). GSH and TSH_2_ can non-enzymatically reduce ovothiol disulfide (OSSO) to its reduced form (OSH).

### Sensitivity of *OvoA* knockouts to anti-leishmanial drugs and inducers of oxidative and nitrosative stress

As the role of ovothiol A as a peroxide scavenger has been proposed [22,23], the sensitivity of *L .mexicana OvoA* knockouts to H_2_O_2_ was tested using a glucose oxidase assay where H_2_O_2_ and D-glucono-δ-lactone are generated continuously from glucose. The knockout strain did not differ with respect to its sensitivity to this glucose oxidase produced H_2_O_2_ (Table 1).

**Table 1:** IC_50_ of different *L. mexicana* cell lines from exposure glucose oxidase, which produces hydrogen peroxide under glucose-rich conditions.

|  | IC<sub>50</sub> ± S. Dev (μM) (n=3) |  | p-value |
| --- | --- | --- | --- |
| Compound | Wild type <sup>(a)</sup> | Δ*OvoA* <sup>(b)</sup> | (b)/(a) |
|  | L. mexicana Cas9 |  |  |
| Glucose oxidase | 3.39 ± 0.8 | 2.91 ± 0.7 | 1.0 (n.s.) |

Shown are mean IC<sub>50</sub> values and Standard deviation from three independent experiments (n=3). A unpaired Student's t-test comparing mutant and Wild type *L. mexicana* was performed. n.s.= non significant

Ovothiols have also been proposed as to play a role in the protection against nitric oxide and nitrosothiols rather than H_2_O_2_ in kinetoplastids given that trypanothione is better adapted electrochemically to reduce H_2_O_2_ [23] . Nevertheless, *OvoA* mutant promastigotes were not found to be more sensitive to the NO-inducing agents SNAP or DETA/NONOate (Table 2).

**Table 2:** IC_50_ of different *L. mexicana* cell lines from exposure to S-nitrosolthiols.

|  | IC<sub>50</sub> ± S. Dev (mM) (n=3) |  | p-value |
| --- | --- | --- | --- |
| Compound | Wild type <sup>(a)</sup> | Δ*OvoA* <sup>(b)</sup> | (b)/(a) |
|  | L. mexicana Cas9 |  |  |
| DETA/NONOate | 0.38±0.005 | 0.35±0.04 | 0.5 (n.s.) |
| SNAP | 0.13±0.002 | 0.19±0.004 | 0.0351 |

Shown are mean IC<sub>50</sub> values and Standard deviation from three independent experiments (n=3). A unpaired Student's t-test comparing mutant and Wild type *L. mexicana* was performed. n.s.= non significant

We also tested the sensitivity of the knockout cells to a range of anti-leishmanial drugs (Table 3). Minor changes (<1.5 fold) were noted with the currently used drugs antimony, paromomycin and miltefosine. The nitroheterocycles fexinidazole, nifurtimox and benznidazole all have anti-leishmanial effects, but in no case was a fold change of >1.4 fold in sensitivity noted in the knockout cells. The only compound tested for which a difference of > 2 fold was seen was amphotericin B where a 2.1 fold reduction in sensitivity was noted (p = 0.0007, Student’s t-test)

**Table 3:** IC_50_ values of different *L. mexicana* lines exposed to antileishmanial drugs.

|  | IC<sub>50</sub> ± SD (µM) |  | p-value |
| --- | --- | --- | --- |
| Drug/ compound | Wild type <sup>(a)</sup> | Δ *OvoA* <sup>(b)</sup> | (b)/(a) |
| L. mexicana Cas9 |  |  |  |
| PAT | 218±3.5 | 200±0.8 | 0.0002 |
| Paromomycin | 92±1.9 | 110±2.8 | 0.0008 |
| Miltefosine | 10±1.5 | 7.2±1.8 | 0.0503 (n.s.) |
| Amphotericin B | 0.18±0.015 | 0.082±0.004 | 0.0007 |
| Fexinidazole | 2.2±0.2 | 2.7±0.15 | 0.0334 |
| Nifurtimox | 8.4±0.4 | 6.01±0.51 | 0.0078 |
| Benznidazole | 160 ±2.4 | 149 ±3.8 | 0.0034 |

Shown are mean IC<sub>50</sub> values and Standard deviation from three independent experiments (n=3). A unpaired Student's t-test comparing mutant and Wild type *L. mexicana* was performed. n.s.= non significant

### *OvoA* cells retain macrophage infectivity and differentiate to amastigotes

Previous studies have shown that ovothiol A is present in metabolite extracts from *L. mexicana* infected macrophages (Clement Regnault, PhD Thesis, University of Glasgow) and ovothiol was also seen in *L. major* amastigotes, albeit not *L. donovani* [7]. It may, hence, be hypothesised that the loss of the *OvoA* gene could affect the ability of *L. mexicana* to infect and proliferate as amastigotes within macrophages. Bone marrow derived macrophages from C57BL/6 mice were infected with stationary -phase Δ*OvoA* and WT *L. mexicana* for 1 week. As seen in Figure 8, intracellular amastigotes were observed in similar numbers in macrophages infected with either the Δ*OvoA* mutant or the WT cell line, suggesting that *OvoA* was non-essential for infection of macrophages, nor for survival and proliferation within these cells.

**Figure 8:**
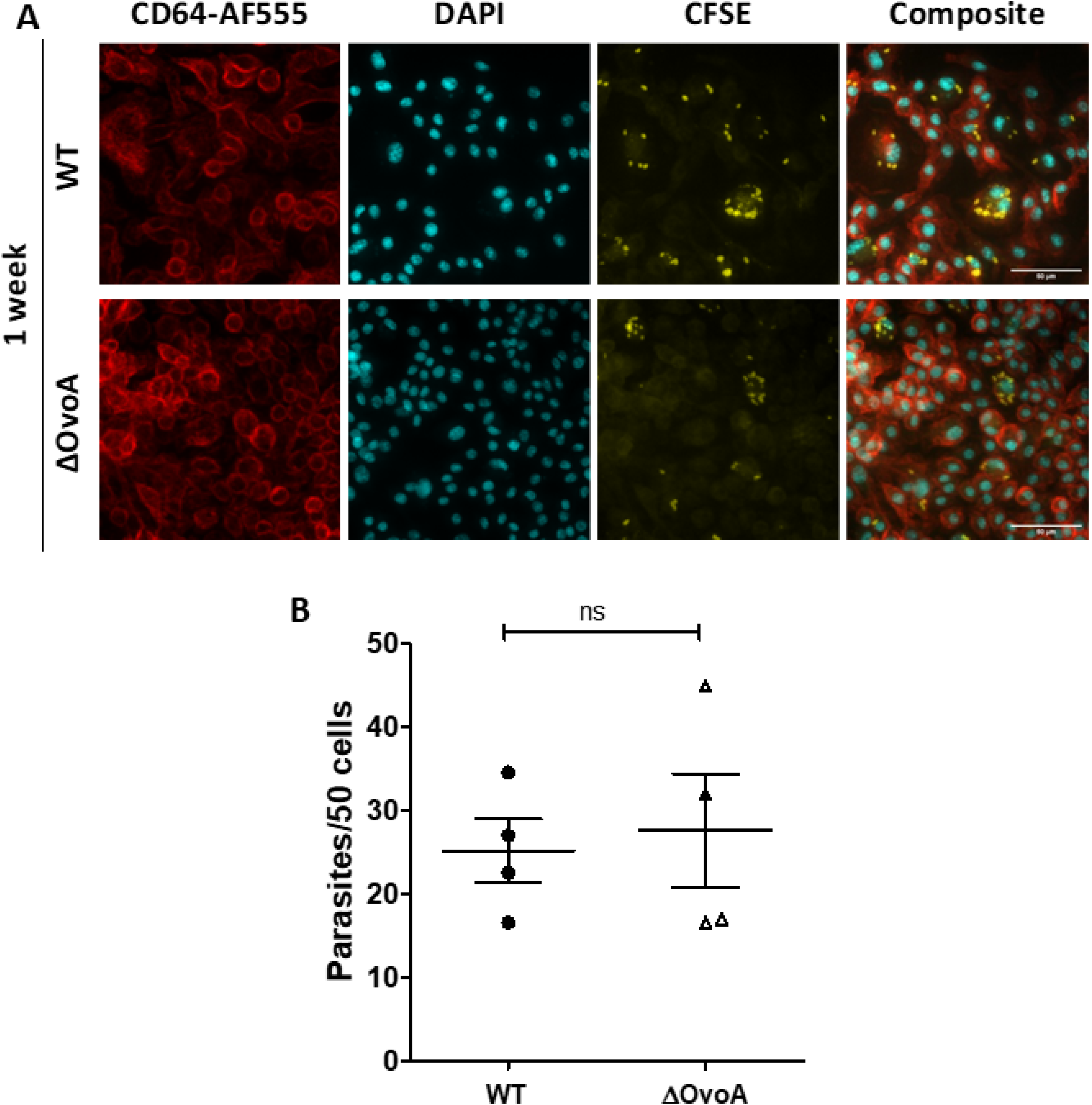
Infection of Bone marrow derived macrophages with Wildype and Δ*OvoA L. mexicana* after 1 week. Representative images of infected macrophages. (B) Quantification of intracellular parasites per 50 macrophages with Fiji Cell counter plugin after 1 week (n=4). (Statistical test: Unpaired t-test. *OvoA*: Ovothiol A; WT: Wild type)

## Discussion

Ovothiol (methyl mercaptohistidine) is a low molecular weight thiol that was originally found in sea urchin eggs [1–4] and later in a variety of other invertebrates including protozoa of the order Kinetoplastida [5–8]. Whilst, in echinoderms, roles in oxidative stress response and also inter-organism communication have been determined, the role of the metabolite in the kinetoplastid parasites has remained elusive, although its low electronegativity and interaction with other thiol-based metabolites, such as trypanothione and glutathione, point to its having roles in cellular redox. More specifically, the fact that it accumulates in metacyclic promastigotes of Leishmania as they prepare for entry into the mammalian host and uptake into macrophages [7], where they need to defend against the oxidative and nitrosative defence mechanisms of those cells, has led to the suggestion that it might play a specific role in detoxifying free radicals [14]. As such, its synthesis has been considered a potentially good target for anti-parasite chemotherapy.

The discovery of an ovothiol synthase enzyme (*OvoA*) [19] and identification of a leishmanial orthologue of the gene encoding the enzyme enabled us to address the function of ovothiol in Leishmania by removing the gene using a CRISPR/cas9 based approach. The knockout cell lines were viable and did not suffer a growth defect as promastigotes when compared to wild type. Moreover, they were able invade and replicate as amastigotes within macrophages. All of this indicates that ovothiol does not play an essential role in the promastigote, or amastigote, form of *L. mexicana* in laboratory conditions.

To confirm that *L. mexicana* ovothiol synthase did, indeed, encode the enzyme required for ovothiol synthesis, we characterised the metabolome of the knockout parasites and compared it to wild type. Two peaks identified as ovothiol A (monomethylated) and ovothiol B (dimethylated) were identified and both were missing from the knockout cells, proving that the enzyme is indeed responsible for the production ovothiol. This approach, knocking out genes that are predicted to encode particular enzymes, followed up with metabolomics analysis offers a route to help identify function of a multitude of genes whose function is currently obscure, especially using the untargeted metabolomics approach where, in principle, any metabolite lost through enzyme-depletion could be found.

The most pronounced other changes to the metabolome in the KO cells also related to thiol metabolism with substantial increases in the abundance in reduced glutathione in the KO cells, and diminished abundance of oxidised glutathione. Reduced trypanothione was also of increased abundance although changes to the oxidised form were not detected using the mass spectrometry platform employed here. This may be because the relative mass spectral signal for the oxidised form was around 100-fold higher than that for the reduced form, and hence large changes in the latter may impact minimally on the signal of the oxidised form. The pronounced increase in reduced forms of both trypanothione and glutathione in the mutant cells, however, would indicate that a significant component of these reduced thiols is involved in maintaining the pool of reduced ovothiol. In the absence of the latter metabolite, then, the reduced forms of the former metabolites increase. Neither glutathione, nor ovothiol are enzymatically reduced in trypanosomatids, instead trypanothione is the central redox metabolite in the system as it is kept in its reduced form by the enzyme trypanothione reductase [24] which uses NADPH to regenerate dihydrotrypanothione from its oxidised disulfide form (Figure 7).

We also compared the metabolome of WT and KO parasites as they progressed through a growth curve from logarithmic to stationary phase. Multiple key metabolic differences accompanied this transformation, for example nucleotides, used in DNA and RNA synthesis were lost in stationary phase while the energy storage polymer, mannogen accumulated. Ovothiol itself accumulates in stationary phase WT *L. mexicana*.

Most anti-leishmanial drugs, including those that are known to induce oxidative stresses [25], were of similar efficacy against ovothiol deficient cells as wild-type. Amphotericin B was an exception, the ovothiol deficient cells being twice as susceptible to this drug. Nitro group containing compounds also had little or no additional activity against the knockout cells, and nor did the nitric oxide generating compounds. It would appear, therefore, that under the conditions used for these assays, ovothiol has little or no effect beyond that afforded by trypanothione/glutathione in protecting the parasites against these key stresses, although the significance of the enhanced activity of amphotericin B is currently unknown, but future work to determine whether pharmacological inhibition of ovothiol production could be used to accentuate the activity of this polyene would be of interest, given the drug is somewhat toxic [26] and reduced dosing would be of practical benefit.

The assays we have performed have aimed to probe potential roles in protection against substantial xenobiotic mediated stress, as it has been presumed that this is the fundamental role of the metabolite. Having failed to identify such a role in these assays, future work should focus on whether specific, more nuanced roles, such is in cell cycle progression or other differentiation states through the life cycle.

## Materials & methods

### CRISPR/cas9 knockout of *OvoA* in *Leishmania mexicana*

Primers for pT-Plasmid and sgRNA template amplification (see Supplementary Table 1) targeting a putative 5-histidylcysteine sulfoxide synthase gene, *OvoA* (LmxM.08_29.0940) were designed using LeishEDIT primer design tool found at: http://leishgedit.net/ [20].

Primers for Diagnostic PCR (see Supplementary Table 2) were designed to target a specific region within the *OvoA* Open reading frame (ORF), as well as its 3’ and 5’ untranslated regions (UTR) using the Primer Blast tool and the *Leishania mexicana* MHOM/GT/2001/U1103 (taxid:929439) reference genome. Output was checked for hairpin loop and primer dimer formation using the OligoAnalyzer Tool (Integrated DNA technologies – www.idtdna.com/). Primers were from Eurofins MWG Synthesis GmbH (Germany).

Blasticidin-S deaminase (c*BSD*) or Puromycin *N*-acetyltransferase (*PAC*) resistant cassettes were amplified from the respective pT-plasmids, pTBlast or pTPuro (kindly provided by Eva Gluenz), using primers containing 5’ and 3’ homologous sequences flanking the *OvoA* target gene. The Master mix for amplification of these genes contained 0.2 mM dNTPs (LT-0224, Thermo Scientific), 2 µM each of gene-specific forward and reverse primers for *OvoA* (LmxM.08_29.0940), 30 ng pT-plasmid either carrying the blasticidin or the puromycin resistance marker and 1 unit HIFI Polymerase (Q5 HF DNA Polymerase, M04915, NEB) and 1× HiFi reaction buffer (05917131103, Roche) supplemented with 1.875 mM MgCl_2_ and 3% DMSO (D8418, Sigma). The total volume was 80 µl. PCR steps were 5 min at 94°C preceding 40 cycles of 30 s at 94°C, 30 s at 65°C, 2 min 15 s at 72°C and a final elongation step for 7 min at 72°C. The product was heat-sterilized at 94°C for 5 min before transfection.

Double stranded sgRNA sequences were created (Supplementary Table 1) to interact with the Cas9 complex and facilitate a double-strand break at the target site. In order to amplify sgRNA templates, 0.2 mM dNTPs (LT-0224, Thermo Scientific), 2 µM of primer G00 (sgRNA scaffold [20]), and 1 unit HiFi Polymerase (Q5 HF DNA Polymerase, M04915, NEB) in 1× HiFi reaction buffer with MgCl_2_ (05917131103, Roche) were mixed with either 5’ sg RNA or 3’ sgRNA forward primer targeting the *OvoA* gene. The total volume was 50 µl. PCR steps were 30s at 98°C preceding 35 cycles of 10 s at 98°C, 30 s at 60°C, 15 s at 72°C.

*Leishmania Cas9* (derived from WHO *L. mexicana* strain MNYC/BZ/62/M379) [20] were transfected in 1x Tb-BSF Buffer using an Amaxa Nucleofector 2b (Lonza) using a single pulse with the X-001 program. 1x 10^7^ parasites were transfected with the PCR products of two sgRNAs targeting the *Leishmania mexicana OvoA* gene and two donor DNA plasmids carrying the blasticidin and puromycin resistance-markers (total combined volume of 100 μl) in an electroporation cuvette in a total volume of 250 μl. After transfection, parasites were transferred into pre-heated HOMEM with 10% FBS and incubated for 16h at 26°C to allow recovery before addition of selection drugs (20 μg/ml puromycin and 5 μg/ml blasticidin).

*L. mexicana OvoA* KO clones were selected using serial 2-fold dilutions starting from 4,000 parasites/well to 0.5 parasites/well were plated on a 96-well plate in HOMEM [27] with 20% FBS and + 20 μg/ml puromycin and 5 μg/ml blasticidin and incubated for up to 14 days at 29°C. Parasites originating from wells calculated as 1 or 0.5 parasites/well were transferred to a T25 cell culture flask in HOMEM with 10% FBS and + 20 μg/ml puromycin and 5 μg/ml blasticidin.

To validate the loss of the *OvoA* gene in the knock out parasite lines, genomic DNA was extracted from 1x10^7^ parasites 12 days after transfection using the DNeasy Blood & Tissue Kit (Qiagen) with non-transfected Cas9 positive and wild type *L. mexicana* serving as negative controls. 2 mM dNTPs, 100 ng of extracted DNA and 2 µM each of forward primer and reverse primer targeting a sequence flanking the *OvoA* gene (P1 – Supplementary Table 2), within the *OvoA* gene (P2 – Supplementary Table 2) or targeting a specific sequence within the puromycin resistant cassette (P3 Supplementary Table 2) were mixed with 1 unit HiFi Polymerase (Q5 HF DNA Polymerase, M04915, NEB) and 1× HiFi reaction buffer with MgCl_2_. The total volume was 50 μl. The PCR steps were 30s of 98°C followed by 35 cycles of 15s of 98°C, 20s at 58°C, 30s at 72°C and a final elongation step for 5 min at 72°C. The absence of the *OvoA* gene was validated by gel electrophoresis on a 1% agarose gel. *L. mexicana Cas9* were propagated in HOMEM (041-94699M, Gibco) with 10% Fetal Bovine Serum (105500, Gibco). Growth medium was supplemented with 2 0μg/ml puromycin and 5 μg/ml blastocidin. The parasite count was determined every 24h for 7 days using a haemocytometer.

### Metabolomics analysis of wild type vs *OvoA* knockout *Leishmania mexicana*

1x 10^8^ stationary phase or log-phase *OvoA* Knockout or WT *Leishmania mexicana* per sample were harvested. Four replicates were collected per condition. Parasites were quenched by cooling to 10°C in a dry ice-ethanol bath and centrifuged at 1,300g for 10 min at 4°C. Supernatant was removed and the cell pellet was washed in ice cold PBS after centrifugation at 1300g for 5 min at 4°C. This step was repeated twice. Sample was centrifuged at 1200g at 4°C for 20 seconds to remove all PBS and the pellet then resuspended in 200 µl of extraction solvent (HPLC grade Chloroform: Methanol:Water-1:3:1) and mixed. Pure extraction solvent served as a blank. Samples were shaken at 4°C. for 1h, before centrifugation at 16.000g for 10 minutes at 4°C. Supernatant was removed and sealed under argon then stored at -80°C until analysis. A pooled sample was generated by mixing supernatant of all samples. Samples were run on LC-MS at Glasgow Polyomics using standard procedures [28]. Briefly this involved separation of metabolites using a ZIC pHILIC column (150 mm × 4.6 mm, 5 μm column, Merck Sequant) before entering an Orbitrap Q-Exactive (Thermo Fisher Scientific) mass spectrometer operating through switching between positive and negative ionization modes. Samples were analysed in four replicates and data were processed picking peaks using mzmatch software [29] then analysed using the Ideom package [30] and Metaboanalyst suite of tools [31]. A cocktail of around 200 metabolites is included in the Glasgow Polyomics workflow in order to provide additional confidence in assigning identities to a number of common metabolites. In the case of trypanothione we obtained a purified sample from Professor Luise Krauth-Seigel (Heidelberg) which was used to provide retention time and mass data to assist in assigning a high level of confidence to trypanothione in the extracted samples (both reduced and oxidised forms were present in the standard).

Metabolomics data have been deposited to the EMBL-EBI MetaboLights database with the identifier MTBLS8790. The complete dataset can be accessed here: www.ebi.ac.uk/metabolights/MTBLS8790.

### Sensitivity of *OvoA* knockout *Leishmania mexicana* to anti-leishmanial drugs and stress inducing agents

Wild type and *OvoA* knockout *L. mexicana* Cas9 parasites were routinely cultured in HOMEM medium (Gibco) supplemented with 10% heat-inactivated fetal bovine serum (Gibco) at 27°C. Antileishmanial drugs (paromomycin, miltefosine, amphotericinB, potassium antimony tartrate, fexinadazole, benznidazole, all from Sigma-Aldrich) were tested against the promastigote stage of wild type and *OvoA* knockout *L. mexicana* Cas9 parasites, measuring IC_50_ values using the Alamar blue assay [32].

In addition, sensitivity to hydrogen peroxide was measured by a glucose oxidase assay [33] (GO from Type VII, Aspergillus Niger, G2133-10KU, solubilised in 1M Potassium phosphate buffer, pH, 6.5), SNAP (N3398, Sigma). Sensitivity to NO-inducing agents S-nitroso-N-acetyl penicillamine (SNAP) (N3398, Sigma) and Z)-1-[2-(2-Aminoethyl)-N-(2-ammonioethyl)amino]diazen-1-ium-1,2-diolate (DETA/NONOate) (ALX-430-014-M005, Enzo Life sciences) involved plating cells in 96-well plates at a final density of 1x10^6^ cells/ml, containing two-fold serial dilutions of each compound and DETA/NONOate (ALX-430-014-M005, Enzo Life sciences), in triplicate. After 72 h of incubation at 27°C, 20 μl of a 0.49 mM resazurin (Sigma-Aldrich) solution was added, and the cultures were incubated for another 48 h at 27°C. Finally, fluorescence was measured at λexcitation = 544 nm and λemission = 590 nm with FLUOstar Optima Microplate Fluorometer (BMG LABTECH GmbH), and IC_50_ values were determined by nonlinear regression from the sigmoidal dose-inhibition curve in GraphPad Prism 5 software (version 5.03).

### Macrophage infection assays

Macrophages were from bone marrow extracts of femurs and tibias from C57BL/6 mice . Bone marrow was removed from bones by flushing with Hank’s Buffered Saline (HBSS, 24020-91, Gibco-Life) and broken down into a single cell suspension with a 25-gauge needle and syringe. The cell suspension was centrifuged at 400 x g for 5 min prior to lysing red blood cells with 1 ml of red blood cell solution for 2 min (Cat. no. 00-4333, eBiosciences). Cells were immediately washed in 10 ml HBSS and pelleted at 400 x g for 5 min at 4°C, then resuspended in 5 ml of HBSS and counted (using a hemacytometer using Trypan Blue dye exclusion, to determine the viable cell concentration). 6x10^6^ cells were plated onto standard sterile 90 mm petri dishes (Sterilin™, Thermo Fisher Scientific) with 10 ml of complete RMPI with 1% Penicillin and Streptomycin and 10% L929 supernatant for infection assay experiments. Cells were incubated at 37° C in a 5% CO_2_ incubator for 8-9 days until cells were fully adhered. Media was changed every 2-3 days. Bone marrow derived macrophages (BMDMs) were harvested, counted and replated at 80,000 cell/well on a 8-well microscope chamber slide (Nunc™ Lab-Tek II ™ Chamber Slide System, Cat# 154534PK Thermofisher Scientific) to prepare for infection experiments.

BMDMs in the monolayer, were infected with stationary phase WT or *OvoA* knock out *L. mexicana*. Promastigote cultures were counted using a hemocytometer and resuspended in DPBS after washing twice (also in DPBS) then labelled with carboxyfluorescein succinimidyl ester (CFSE, CellTrace™ CFSE Cell Proliferation Kit, Cat# C34554,Thermofisher Scientific) in DPBS (1:1000). Following staining for 10 minutes at 29°C in the dark, promastigotes were washed with DPBS, pelleted (800g, 10min) and resuspended in complete RMPI with 1% Penicillin/streptomycin (100 units penicillin and 0.1 mg/ml streptomycin) (Penicillin-Streptomycin 100x, Cat#. P4333, Sigma-Aldrich). Subsequently,1.6x10^5^ parasites/well (MOI: 2:1) were added to the macrophage cultures and incubated at 34°C with 5% CO_2_ for 6 h before being washed twice in DPBS to remove extracellular promastigotes and incubated in cRMPI + 10% L929 at 34°C with 5% CO_2_ for 1 week until further processing for immunofluorescent staining. Media was replaced every 2-3 days.

Media was removed from wells of microscope chambers slides and wells were carefully rinsed with 1% BSA in DPBS. Cells were fixed for 20 minutes with 4% paraformaldehyde, followed by two 5 minute-washes in 1% BSA in PBS on a shaker. Cells were stained with anti-CD64 (CD64 Recombinant Rabbit Monoclonal Antibody, Thermofisher, **#** MA5-29706) at a dilution of 1:400 in 1% BSA in PBS and incubated at 4°C overnight in the dark overnight. The next day, antibodies were removed by two washes as described, followed by staining with secondary anti-Rabbit antibodies (1:500) conjugated to Alexa Fluor 555 (Goat anti-Rabbit IgG, Superclonal Recombinant Secondary Antibody, Alexa Fluor 555, Thermofisher, # A27039) for 30 minutes on a shaker at Room temperature. Cells were carefully washed twice in PBS for 5 minutes, followed by 10 minute incubation with 5 μg/ml of DAPI in PBS. Cells were washed twice in 1% BSA in PBS before the chamber was carefully removed. Slides were sealed with Vector Shield and a coverslip and stored overnight at 4 °C to dry, then analysed on an Inverted Zeiss SpinningDisk microscope. Parasite counts were obtained using the Cell counter plugin in Fiji Image J.

## Supporting information captions

Supplementary Table 1: Oligos for CRISPR replacement cassette and sgRNA amplification

Supplementary Table 2: Oligos for diagnostic PCRs to validate *OvoA* Knockout

Supplementary Table 3: Oligos for diagnostic PCRs to validate *OvoA* Knockout

**Supplementary Figure S1a: Schematic representation of target gene locus before and after successful CRISPR Cas9-mediated Knockout with expected product sizes of validation reactions by PCR.**

**Supplementary Figure S1b: Gel Validation of Crispr Knock out of LmxM.08_29.0940 (*OvoA*) of three clones (GLG922, GLG923, GLG950) from two independent transfections.** Shown are Polymerase chain reaction (PCR) products of Knock out clones GLG922, GLG923, GLG950, Wild type and negative control of PCRs with (A) Primer pair P1 flanking the target gene locus, (B) Primer pair P2 targeting the Puromycin resistance cassette, Primer pair P3 binding specifically within the target gene (C) and a positive control amplifying the PF16 gene (C). GLG922 and GLG923 show successful insertion of puromycin and blasticidin resistance cassettes (A) alongside removal of the OvoA Gene (C). GLG950 demonstrates removal of OvoA Gene via PCR (C) but no insertion of puromycin and blasticidin resistance cassettes (A). Clone GLG923 was used in subsequent experiments.

**Supplementary S2: Heatmap of top 100 siginificantly changed meatbolites in log-phase (A) and stationary phase (B) ΔOVOA and Wildtype *L. mexicana.*** Heatmaps were generated with the metaboanlayst vs5.0 online tool using the Eucladian (Ward) clustering method, and Scaling method set to Autoscaling by features. OVOA: Ovothiol (red) A; WT: Wildtype (green).

**Supplementary Figure S3: Metabolome Sets Enrichment analysis of significantly up-(A) and downregulated metabolites(B) in logarithmic-phase OVOA and WT *L. mexicana.*** Threshold for significance were p. adj. (FDR)<0.05 and Log_2_(FC)>1. Enrichment analysis was performed using the inbuilt Enrichment analysis tool from metaboanalyst vs.5.0. The length of the bars in the barchart, signify the Enrichment ratio.The Colorshading signifies the significance of the Enrichment pathway selection (Red: Most significant< White: Least significant)

**Supplementary Figure S4: Metabolome Sets Enrichment analysis of significantly up-(A) and downregulated metabolites (B) in stationary-phase OVOA and WT *L. mexicana.*** Threshold for significance were p. adj. (FDR)<0.05 and Log_2_(FC)>1. Enrichment analysis was performed using the Enrichment analysis tool from metaboanalyst vs.5.0. The length of the bars in the barchart, signify the Enrichment ratio. The Colorshading signifes the significance of the Enrichment Pathway selection (red: Most sinificant, white: least significant).

**Supplementary Figure S5: Peak intensities of three different peaks identified as putative trypanothione (A, B, C) and of trypanothione disulfide (D) and corresponding peaks extracted from Ideom (E,F, G,H) in logarithmic and stationary phase ΔOvoA and WT *L. mexicana*.** Each group is composed of four metabolomic samples (n=4). Statistical significance was tested using a One-way ANOVA with Bonferronis Multiple Comparison Test as a post-hoc test (***=p<0.001; *=p<0.05). Chromatographs display the retention time (RT) on the X axis and peak intensity on the Y axis.

**Supplementary Figure S6: Mass, Retention time and Ion mode of putative Trypanothione and trypanothione disulfide peaks displayed in Supplementary Figure S5.**

**Supplementary Figure S7: Extracted Iron chromatography in negative ion mode (A) and positive ion mode (B) of Trypanothione standard (top) and Test sample OvoA KO (bottom).**

## References

1. Turner E, Klevit R, Hopkins PB, Shapiro BM. Ovothiol: a novel thiohistidine compound from sea urchin eggs that confers NAD(P)H-O2 oxidoreductase activity on ovoperoxidase. J Biol Chem. 1986;261(28):13056–63. doi:10.1016/S0021-9258(18)69270-1

2. Turner E, Klevit R, Hager LJ, Shapiro BM. Ovothiols, a family of redox-active mercaptohistidine compounds from marine invertebrate eggs. Biochemistry. 1987;26(13):4028–36. doi: 10.1021/bi00387a043.

3. Castellano I, Seebeck FP. On ovothiol biosynthesis and biological roles: from life in the ocean to therapeutic potential. Nat Prod Rep. 2018;35(12):1241–1250. doi: 10.1039/c8np00045j.

4. Cordell GA, Lamahewage SNS.Ergothioneine, Ovothiol A, and selenoneine-histidine-derived, biologically significant, trace global alkaloids. Molecules. 2022;27(9):2673. doi: 10.3390/molecules27092673.

5. Steenkamp DJ, Spies HS.Identification of a major low-molecular-mass thiol of the trypanosomatid *Crithidia fasciculata* as ovothiol A. Facile isolation and structural analysis of the bimane derivative. Eur J Biochem. 1994;223(1):43–50. doi: 10.1111/j.1432-1033.1994.tb18964.x.

6. Spies HS, Steenkamp DJ.Thiols of intracellular pathogens. Identification of ovothiol A in *Leishmania donovani* and structural analysis of a novel thiol from *Mycobacterium bovis*. Eur J Biochem. 1994;224(1):203–13. doi: 10.1111/j.1432-1033.1994.tb20013.x.

7. Ariyanayagam MR, Fairlamb AH. Ovothiol and trypanothione as antioxidants in trypanosomatids. Mol Biochem Parasitol. 2001;115(2):189–98. doi: 10.1016/s0166-6851(01)00285-7.

8. Steenkamp DJ. Trypanosomal antioxidants and emerging aspects of redox regulation in the trypanosomatids. Antioxid Redox Signal. 2002;4(1):105–21. doi: 10.1089/152308602753625906.

9. Barrett MP, Burchmore RJ, Stich A, Lazzari JO, Frasch AC, Cazzulo JJ, Krishna S. The trypanosomiases. Lancet. 2003;362(9394):1469–80. doi: 10.1016/S0140-6736(03)14694-6.

10. Burza S, Croft SL, Boelaert M. Leishmaniasis. Lancet. 2018;392(10151):951–970. doi: 10.1016/S0140-6736(18)31204-2.

11. Fairlamb AH, Blackburn P, Ulrich P, Chait BT, Cerami A.Trypanothione: a novel bis(glutathionyl)spermidine cofactor for glutathione reductase in trypanosomatids. Science. 1985;227(4693):1485-7. doi: 10.1126/science.3883489.

12. Fairlamb AH, Cerami A. Metabolism and functions of trypanothione in the Kinetoplastida. Annu Rev Microbiol. 1992;46:695–729. doi: 10.1146/annurev.mi.46.100192.003403.

13. Leroux AE, Krauth-Siegel RL. Thiol redox biology of trypanosomatids and potential targets for chemotherapy. Mol Biochem Parasitol. 2016;206(1-2):67–74. doi: 10.1016/j.molbiopara.2015.11.003.

14. Vogt RN, Steenkamp DJ. The metabolism of S-nitrosothiols in the trypanosomatids: the role of ovothiol A and trypanothione. Biochem J. 2003;371(Pt 1):49–59. doi: 10.1042/BJ20021649.

15. Kaye P, Scott P. Leishmaniasis: complexity at the host-pathogen interface. Nat Rev Microbiol. 2011;9(8):604–15. doi: 10.1038/nrmicro2608.

16. Desjardins M, Descoteaux A. Survival strategies of Leishmania donovani in mammalian host macrophages. Res Immunol. 1998;149(7-8):689–92. doi: 10.1016/s0923-2494(99)80040-6.

17. Ferreira C, Estaquier J, Silvestre R. Immune-metabolic interactions between Leishmania and macrophage host. Curr Opin Microbiol. 2021;63:231–237. doi: 10.1016/j.mib.2021.07.012.

18. Braunshausen A, Seebeck FP. Identification and characterization of the first ovothiol biosynthetic enzyme. J Am Chem Soc. 2011;133(6):1757–9. doi: 10.1021/ja109378e.

19. Goncharenko KV, Vit A, Blankenfeldt W, Seebeck FP. Structure of the sulfoxide synthase EgtB from the ergothioneine biosynthetic pathway. Angew Chem Int Ed Engl. 2015 Feb 23;54(9):2821–4. doi: 10.1002/anie.201410045.

20. Beneke T, Madden R, Makin L, Valli J, Sunter J, Gluenz E. CRISPR Cas9 high-throughput genome editing toolkit for kinetoplastids. .R Soc Open Sci. 2017;4(5):170095. doi: 10.1098/rsos.170095.

21. Ralton JE, Sernee MF, McConville MJ. Evolution and function of carbohydrate reserve biosynthesis in parasitic protists. Trends Parasitol. 2021;37(11):988–1001. doi: 10.1016/j.pt.2021.06.005. PMID: 34266735

22. Bailly F, Zoete V, Vamecq J, Catteau JP, Bernier JL. Antioxidant actions of ovothiol-derived 4-mercaptoimidazoles: glutathione peroxidase activity and protection against peroxynitrite-induced damage. FEBS Lett. 2000;486(1):19–22. doi: 10.1016/s0014-5793(00)02234-1.

23. Osik NA., Zelentsova EA, Tsentalovich YP. Kinetic studies of antioxidant properties of ovothiol A, Antioxidants. MDPI, 2021; 10(9). doi:10.3390/ANTIOX10091470/S1.

24. Shames SL, Fairlamb AH, Cerami A, Walsh CT. Purification and characterization of trypanothione reductase from *Crithidia fasciculata*, a newly discovered member of the family of disulfide-containing flavoprotein reductases. Biochemistry 1986; 17;25(12):3519–26. doi: 10.1021/bi00360a007.

25. Moreira W, Leprohon P, Ouellette M. Tolerance to drug-induced cell death favours the acquisition of multidrug resistance in Leishmania. Cell Death Dis. 011;2(9):e201. doi: 10.1038/cddis.2011.83.

26. Cavell G. The Problem with Amphotericin. Clin Drug Investig. 2020;40(8):687–693. doi: 10.1007/s40261-020-00924-4.

27. Berens RL, Brun R, Krassner SM. A simple monophasic medium for axenic culture of hemoflagellates. J Parasitol. 1976;62(3):360–5.

28. Kovářová J, Pountain AW, Wildridge D, Weidt S, Bringaud F, Burchmore RJS, et al.. Deletion of transketolase triggers a stringent metabolic response in promastigotes and loss of virulence in amastigotes of Leishmania mexicana. PLoS Pathog. 2018;14(3):e1006953. doi: 10.1371/journal.ppat.1006953.

29. Scheltema RA, Jankevics A, Jansen RC, Swertz MA, Breitling R. PeakML/mzMatch: a file format, Java library, R library, and tool-chain for mass spectrometry data analysis. Anal Chem. 2011;83(7):2786–93. doi: 10.1021/ac2000994.

30. Creek DJ, Jankevics A, Burgess KE, Breitling R, Barrett MP. Ideom: an Excel interface for analysis of LC-MS-based metabolomics data. Bioinformatics. 2012;28(7):1048–9. doi:10.1093/bioinformatics/bts069.

31. Chong J, Wishart DS, Xia J. sing MetaboAnalyst 4.0 for Comprehensive and Integrative Metabolomics Data Analysis. Curr Protoc Bioinformatics. 2019;68(1):e86. doi: 10.1002/cpbi.86.

32. Mikus J, Steverding D.A simple colorimetric method to screen drug cytotoxicity against Leishmania using the dye Alamar Blue Parasitol Int. 2000;48(3):265–9. doi: 10.1016/s1383-5769(99)00020-3.

33. Channon JY, Blackwell, JM. A study of the sensitivity of Leishmania donovani promastigotes and amastigotes to hydrogen peroxide. I. Differences in sensitivity correlate with parasite-mediated removal of hydrogen peroxide, Parasitology. 1985; 91 (Pt 2): 197–206. doi: 10.1017/S0031182000057309.

